# Differential stage-specific mortality as a mechanism for diversification

**DOI:** 10.1101/2022.07.01.498500

**Authors:** P. Catalina Chaparro-Pedraza

**Affiliations:** Swiss Federal Institute of Aquatic Science and Technology (EAWAG), Dübendorf, Switzerland

## Abstract

Individual variability in mortality is widespread in nature. The general rule is that larger organisms have a greater chance of survival than smaller conspecifics. There is growing evidence that differential mortality between developmental stages has important consequences for the ecology and evolution of populations and communities. However, we know little about how it can influence diversification. Using an eco-evolutionary model of diversification that considers individual variability in mortality, we show that commonly observed differences in mortality between juveniles and adults facilitate ecological diversification. We find that juvenile-biased mortality reduces the threshold of minimum resource productivity required for diversification. Diversification is hence less restricted when mortality is more biased towards juveniles than when all individuals experience the same mortality rate. This is because, by altering the population composition, juvenile-biased mortality increases the strength of intraspecific competition. Strong intraspecific competition in turn induces frequency-dependent selection, which drives ecological diversification. Our results demonstrate the strong influence that differential mortality between developmental stages has on diversification, and highlight the need for integrating developmental processes into the theoretical framework of diversification.

## Introduction

Ecological diversification is thought to have given rise to much of the diversity of life on earth^1,2^. This process, in which a single ancestral population diversify into ecologically different species that exploit a variety of niches^2^, has produced several diverse adaptive radiations including Darwin’s finches^3^, Caribbean anole lizards^4^, and cichlid fishes^5^. To understand the origin of biodiversity, it is therefore fundamental to identify the factors facilitating or constraining ecological diversification.

Intraspecific competition for food resources has been extensively investigated as a mechanism for ecological diversification^6,7^. Strong competition can induce frequency-dependent selection causing the fitness landscape to change in response to evolutionary change. As a consequence, adaptive evolution can drive phenotypic traits towards a local minimum of the fitness landscape^8^. At this point, intermediate phenotypes have a fitness disadvantage compared with more extreme phenotypes, causing disruptive selection and thus ecological diversification^9^. Any factor modulating the strength of competition is thus essential for determining whether ecological diversification by this mechanism takes place or not. For instance, increased mortality weakens resource competition, resulting in frequency-independent selection and thus no diversification^10^.

Theory on ecological diversification is founded on the assumption that individuals within a population do not differ in their developmental stage^11,12^ (but see^13^). Yet, during development, individuals experience changes in their ecological role, and these changes can have large impacts on the ecology and evolution of populations and communities^14–18^. Mortality, for instance, varies considerably across developmental stages and this variation can stabilize food webs^17^, alter functional community composition^14^, and cause evolutionary-driven regime shifts^16^. Generally, smaller or younger individuals of diverse fish ^19–21^, amphibian ^22–24^, reptile^25–27^ and invertebrate species^28–30^ are more vulnerable to predation, less resistant to starvation, and less tolerant to environmental extremes than larger ones. Differential mortality between life stages is therefore ubiquitous in nature and has the potential to alter the strength of competition and thus the outcome of diversification. Yet, the effect of differential mortality between life stages on ecological diversification remains to be investigated.

The aim of this study is to understand how differential mortality between life stages affect ecological diversification. We address this question using an eco-evolutionary model employed in the past to investigate ecological diversification driven by intraspecific competition for food resources^10^, and extending it to include demographic dynamics of two different life stages (figure 1A). In the model, an ecological trait that determines resource use can evolve in response to natural selection. The fitness landscape is dynamical: fitness increases with food availability, which in turn is altered by the density of individuals and their trait. Diversification occurs through a process of evolutionary branching, whereby a population evolves to a trait value where it undergoes disruptive selection and splits up into two phenotypically diverging lineages^31,32^. We examine how differential mortality between life stages affect the evolutionary dynamics of a population colonizing a habitat with two food resources, and investigate under which conditions selection favors diversification.

**Figure 1.**
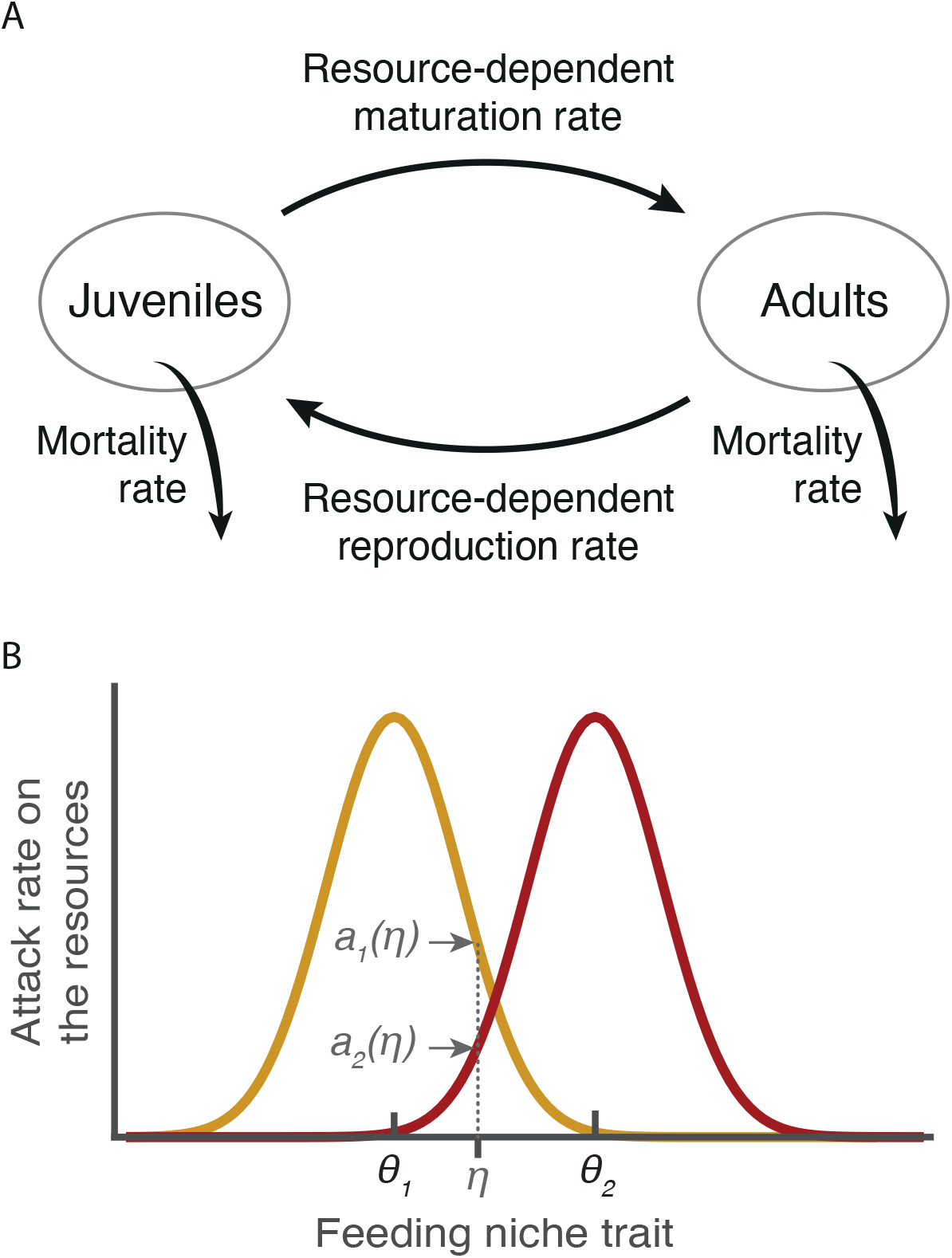
Model structure. A) The model considers two life stages: juveniles and adults. Newborn individuals start their life as juveniles. During the juvenile stage, individuals use their food intake to mature. Once mature, individulas in the adult stage use their food intake for reproduction. B) The model also considers two food resources that have an optimal feeding niche trait *θ*_*1*_ and *θ*_*2*_ to be consumed. The feeding niche trait determines the resource use (e.g. the maximum gape in fish or reptiles, or the bill size in birds, determines the size of the food particules that they can ingest). The gaussian curves describe how the attack rate on each resource, *a*_*1*_*(η)* and *a*_*2*_*(η)*, varies with the feeding niche trait *η*. Individuals are therefore characterized by their feeding niche trait and their developmental stage. The feeding niche trait determines their food intake and thus their maturation and reproduction rate, whereas the developmental stage determines the mortality rate. The model can be readily extended to consider several food resources, differential feeding rates between life stages, and metabolic maintenance costs (see Supporting Information 1).

## Methods

We formulate an eco-evolutionary model of ecological diversification driven by competition for multiple resources. Similar models have been used to investigate the effect of ecological interactions on diversification^10^. We extend these models to include life stages differences, in particular differences in mortality.

### Food resources and feeding

The model considers two food resources with density *F*_*i*_ (*i* = 1,2), but it can be readily extended to consider more resources (Supplementary Information 1). In the absence of consumers, resource density dynamics follow 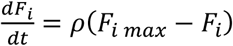, where *ρ* and *F*_*i max*_ are the renewal rate and the carrying capacity of the *i*th resource, respectively. The total productivity is the sum of the carrying capacities of the resources *P* = ∑_*i*_ *F*_*i max*_. We assume *F*_1 *max*_ = *F*_2 *max*_ = *P*/2. However, deviations from this assumption are investigated in Supplementary information (figure S2).

We consider *m* emerging ecomorphs (*j* = 1, …, *m*) characterized by its feeding niche trait *η*_*j*_, which determines their resource use. The attack rate of an ecomorph with feeding niche trait *η*_*j*_ on the *i*th resource, *a*_*i*_(*η*), equals the maximum attack rate *α* when its feeding niche trait *η*_*j*_ equals *θ*_*i*_, and decreases in a Gaussian manner as *η*_*j*_ moves away from *θ*_*i*_, that is *a*_*i*_(*η*_*j*_) = *α* exp [−(*η*_*j*_− *θ*_i_)^2^/(2*τ*^2^)] (figure 1B). In this expression, *τ* determines the width of the Gaussian function and hence, when it is small, an ecomorph must be highly specialized to successfully attack the resource *i*. Individuals feed at a rate *a*_*i*_(*η*_*j*_)*F*_*i*_ on the *i*th resource, and thus deplete it at this rate.

### Life stages

The model considers an ecomorph population to be composed of juvenile and adult individuals. Juveniles mature (i.e. pass to the adult stage) and adults reproduce at a rate proportional to their feeding rate. Individuals can die due to background mortality *δ*_*b*_ and stage-specific mortality. To implement differential mortality between life stages, we follow previous work^17^ and thus assume stage-specific mortality to be equal to *ϕδ*_*s*_ and (2 − *ϕ*)*δ*_*s*_ for juvenile and adults, respectively. In this expression *δ*_*s*_ is the maximum stage-specific mortality and *ϕ* is the differential mortality factor. This factor, ranging between 0 and 2, determines whether mortality is biased towards juveniles (*ϕ* > 1), adults (*ϕ* < 1), or equal for all individuals (*ϕ* = 1). Assuming mortality to be equal for all individuals is equivalent to assume that mortality is independent of the developmental stage, as it has been implicitly assumed by existing theory on ecological diversification, which neglects the existence of distinct stages.

### Ecological dynamics

We model the ecological dynamics considering the densities of juveniles *J*_*j*_ and adults *A*_*j*_ of the *j*th ecomorph, as well as of food resources *F*_*i*_. Based on the assumptions described above, the ecological dynamics follow

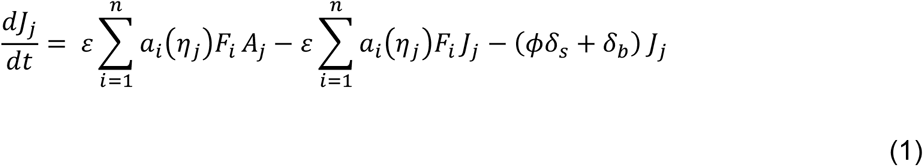

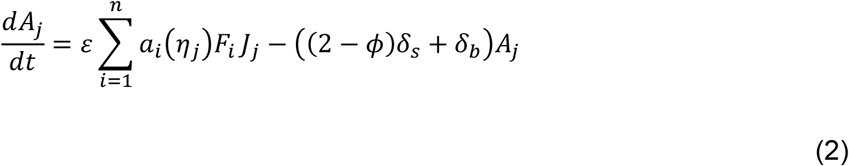

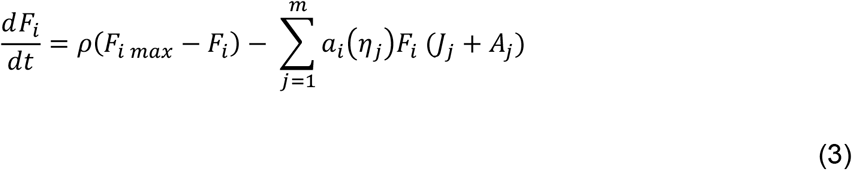

where, *ε* is the efficiency with which food is converted to biomass.

### Evolutionary dynamics

We model the evolution of *m* ecomorphs with trait values *η* = (*η*_1_, …, *η*_*m*_) using the adaptive dynamics framework^33^. This framework assumes that evolution occurs via small, infrequent mutational steps, and thus that the system reaches the ecological equilibrium in between two mutations. Under this assumption, the rate of change in the feeding niche trait of the *j*th ecomorph population can be approximated by the canonical equation^34^:

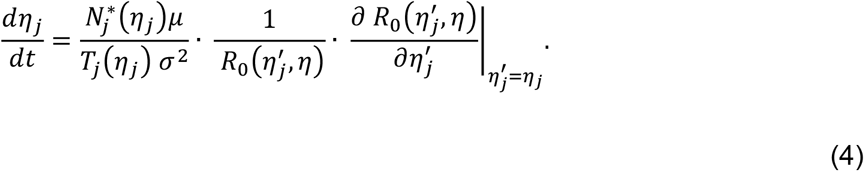

In this expression, *μ* is the mutation rate per birth event, *σ*^2^ is the variance of the trait offspring distribution, *T*_*j*_ (*η*_*j*_) is the expected lifespan of the *j*th resident ecomorph and 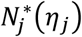 its density, i.e. 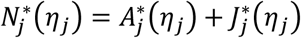. 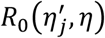 is the lifetime reproductive output of a mutant with trait 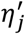 in the environment of the resident ecomophs with trait values *η*. The derivation of 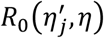 can be found in Supplementary information 2. The term 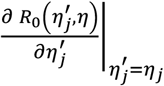 is proportional to the selection gradient.

Directional selection ceases when the selection gradient vanishes. At this point, selection can be stabilizing if the trait value at the equilibrium corresponds to a fitness maximum, or disruptive if it corresponds to a fitness minimum. The curvature of the fitness function therefore determines whether evolution halts or an ecomorph splits into two different ecomorphs. The calculation of the curvature of the fitness function is described in Supplementary information 2.

### Model analysis

We investigate how the difference in mortality between life stages affect diversification when one population colonizes an environment with two different food resources (an extended version of the model that considers more resources can be found in Supplementary information 1). We simulate the evolutionary dynamics when the mortality rate of juveniles and adults is equal (*ϕ* = 1) and when mortality is biased towards juveniles (*ϕ* = 1.6) (figure 2A, 2B). Throughout the simulation we follow the number of ecomorphs, their feeding niche trait and the densities of juveniles, adults and food resources, which we use to calculate the fitness gradient (eq. SI1.5, Supplementary information 1). With the fitness gradient and the curvature of the fitness landscape (eq. SI1.6, Supplementary information 1), we investigate how the difference in mortality between life stages (figure 2C) and the productivity (figure 4) influence diversification. We furthermore investigate the effect of the difference in mortality between life stages on the population and resource densities in the equilibrium at different fixed trait values using eq. 1-3 (figure 3).

**Figure 2.**
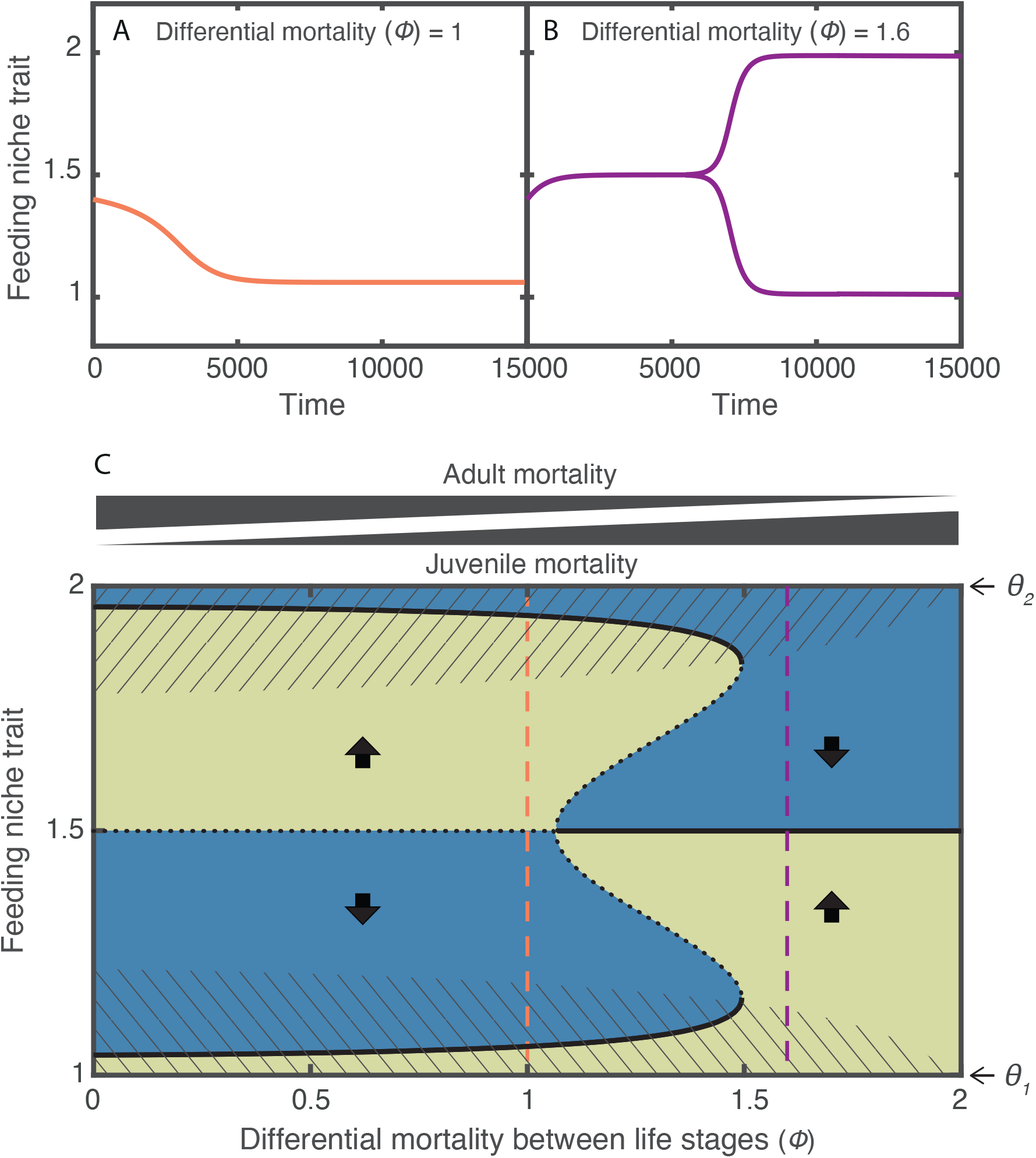
Juvenile-biased mortality promotes diversification. Evolutionary dynamics of a population colonizing an environment with two different food resources and whose individuals experience A) equal mortality in the juvenile and the adult stage, and B) juvenile-biased mortality. C) Fitness gradient of this population as a function of differential mortality between life stages. Diversification occurs only at high values of the differential mortality factor because the trait of the population evolves towards the evolutionary equilibrium that corresponds to a fitness minimum, i.e. where the trait equals 1.5 (arrows indicate the direction of selection). On the contrary, when the differential mortality factor is low, the population evolves toward the vecinity of either of the optimal traits to feed on the resources, i.e. in the neighborhood of *θ*_*1*_ or *θ*_*2*_, depending on the initial trait value of the population. These trait values correspond to fitness maxima, therefore diversification does not occur. Background color indicates the selection gradient (yellow when positive and blue when negative). Evolutionary equilibriums are indicated by black thick lines (solid when they are attractors and dotted when they are repellers). Evolutionary equilibriums in the striped regions are fitness maxima, whereas outside these regions they are fitness minima. Vertical color lines correspond to the fitness gradient experienced by the populations simulated in A (pink) and in B (purple, until time 5500 when the population splits in two). Parameter values as in table 1.

**Figure 3.**
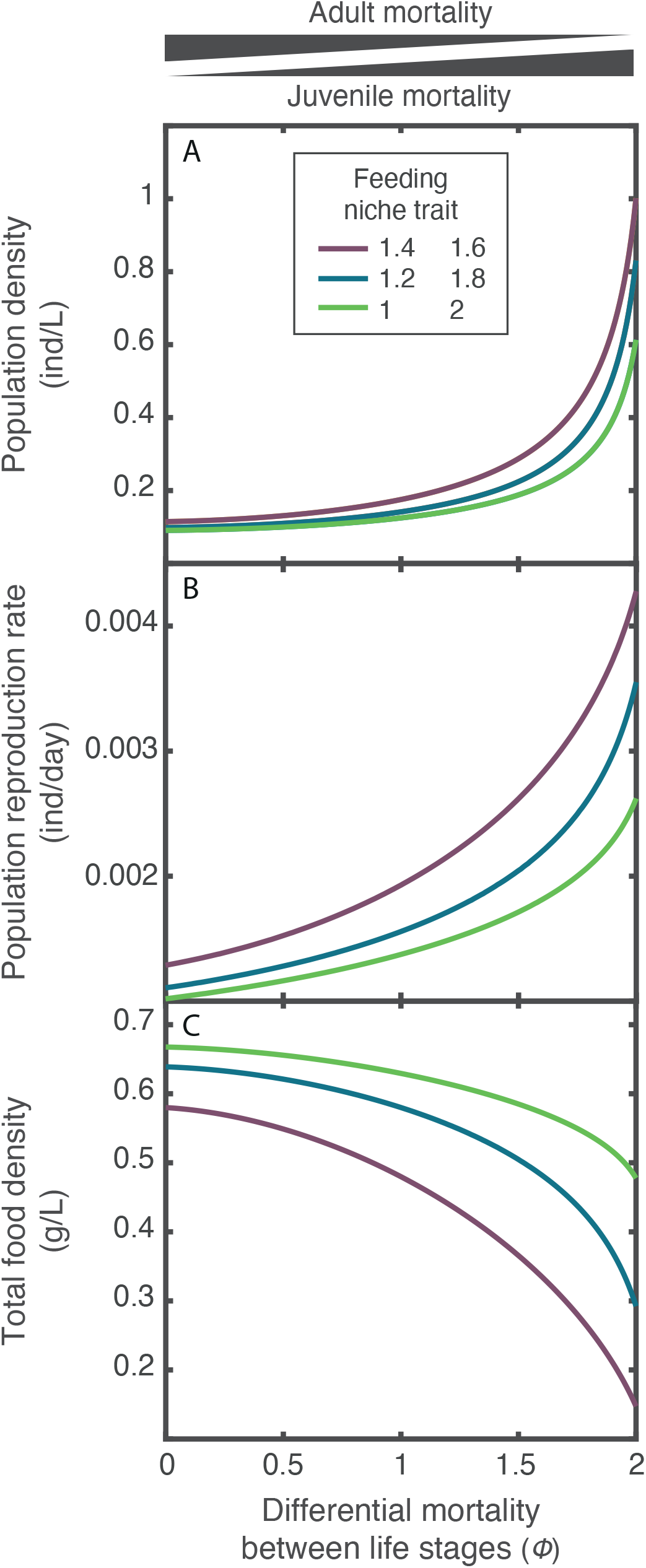
Juvenile-biased mortality increases population density. A) Population density, B) population reproduction rate and C) total food density in the ecological equilibrium as a function of the differential mortality between life stages (*ϕ*). The population density is the sum of the juvenile and adult density. The total food density is the sum of the densities of the two food resources. The system is symmetric around the trait value in between the optimal traits to feed on the resources, i.e. 1.5. Hence, the densities are equal for the trait values that are equidistant from this trait value (e.g. 1.4 and 1.6). No evolutionary dynamics are considered, the densities in the ecological equilibrium are thus computed using eq. 1-3. Parameter values as in table 1.

**Figure 4.**
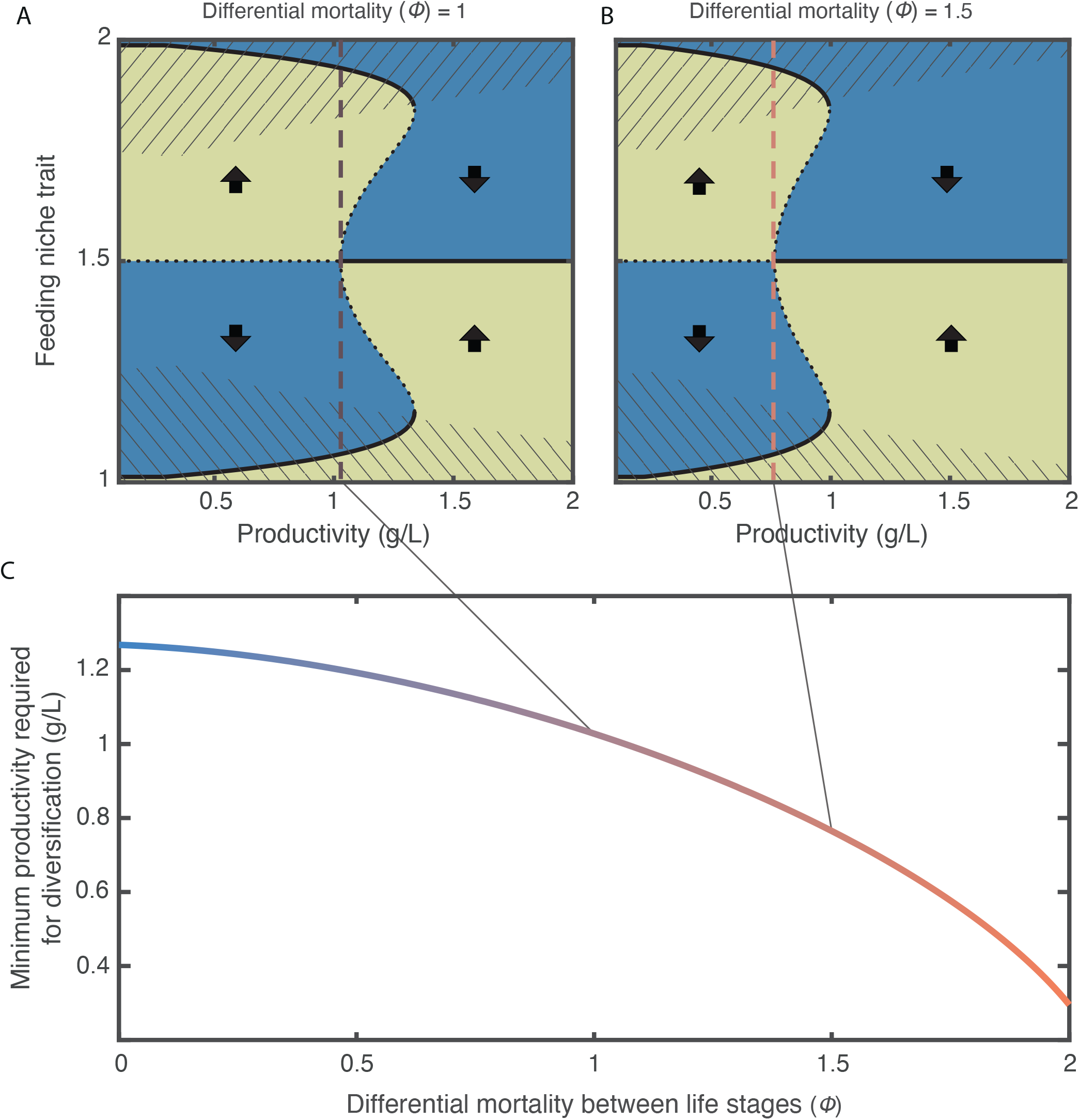
Juvenile-biased mortality facilitates diversification. Fitness gradient of a population colonizing an environment with two different food resources and whose individuals experience A) equal mortality in the juvenile and the adult stage, and B) juvenile-biased mortality. C) Minimum productivity required for diversification as a function of the factor of differential mortality between life stages. Because diversification occurs only when the population evolves towards the evolutionary equilibrium that corresponds to a fitness minimum (i.e. where the trait equals 1.5), the minimal productivity for diversification is that at which this equilibrium changes from a repeller (at low productivity) to an attractor (at high productivity) of the evolutionary dynamics. In A and B, background color indicates the selection gradient (yellow when positive and blue when negative). Evolutionary equilibriums are indicated by black thick lines (solid when they are attractors and dotted when they are repellers). Evolutionary equilibriums in the striped regions are fitness maxima, whereas outside these regions they are fitness minima. Parameter values as in table 1.

## Results

### Juvenile-biased mortality enables ecological diversification

The evolutionary dynamics of a population colonizing an environment with two food resources shows that when juveniles and adults experience equal mortality, the feeding niche trait evolves towards the vicinity of one of the optima to feed on the resources (figure 2A). At this point, diversification cannot occur because selection is balancing. Conversely, when juveniles experience higher mortality than adults, the feeding niche trait evolves to the value in between the two optima (trait value 1.5). At this point, selection is disruptive, hence the more extreme phenotypes are selected for and the population undergoes a diversification event. After diversification, the traits of the two resulting populations diverge. Each population evolves towards one of the trait values that is optimal to feed on the resources (figure 2B).

**Table 1.** Variables and parameters

| Variable or parameter | Symbol | Value | Units |
| --- | --- | --- | --- |
| Variables |  |  |  |
| Feeding niche trait | $\eta$ | Evolving trait | - |
| Juvenile density | $J_j$ | - | $L^{-1}$ |
| Adult density | $A_j$ | - | $L^{-1}$ |
| Food resource density | $F_i$ | - | $g L^{-1}$ |
| Parameters of the habitat |  |  |  |
| Renewal rate of resources† | $\rho$ | 0.01 | $(\text{unit of time})^{-1}$ |
| Total productivity of the habitat | $P$ | 1 | $g L^{-1}$ |
| Ratio of maximum resource density between the resources ($F_{1\max}/F_{2\max}$) | $r$ | 1 (by default)<br>varied in fig. S2 | - |
| Optimum to feed on resource 1 | $\theta_1$ | 1 | - |
| Optimum to feed on resource 2 | $\theta_2$ | 2 | - |
| Demographic parameters |  |  |  |
| Maximum attack rate† | $\alpha$ | 0.2 | $L (\text{unit of time})^{-1}$ |
| Width of the Gaussian curve describing the degree of specialization to successfully attack a food resource | $\tau$ | 1/3 | - |
| Assimilation efficiency | $\varepsilon$ | 0.6 | - |
| Background mortality rate† | $\delta_b$ | 0.001 | $(\text{unit of time})^{-1}$ |
| Maximum stage-specific mortality rate† | $\delta_s$ | 0.01 (by default)<br>varied in fig. S1 | $(\text{unit of time})^{-1}$ |
| Differential mortality factor | $\phi$ | varied (0-2) | - |
| Evolutionary parameters |  |  |  |
| Mutation rate | $\mu$ | $10^{-3}$ (by default)<br>varied in fig. S6 | - |
| Variance of the trait offspring distribution | $\sigma^2$ | 0.01 | - |

†Rates have an arbitrary unit of time (e.g. day)

An analysis of the fitness gradient experienced by a population colonizing an environment with two food resources reveals that differential mortality between life stages alters the stability of the evolutionary equilibrium existing in between the two optima to feed on the resources (feeding niche trait 1.5). This evolutionary equilibrium resides at a local minimum of the fitness landscape, therefore a population with this trait value experiences disruptive selection. When mortality is biased towards adults, this equilibrium is a repeller of the evolutionary dynamics, in other words, evolution drives the feeding niche trait value away from it (left in figure 2C). Conversely, when mortality is biased towards juveniles, this evolutionary equilibrium is an attractor of the evolutionary dynamics, and thus the feeding niche trait evolves towards it (right in figure 2C). Because disruptive selection has a diversifying effect only when a local fitness minimum is an attractor of the evolutionary dynamics^9^, diversification can occur only when mortality is biased towards juveniles.

### Differential mortality between life stages alters the strength of intraspecific competition

Having observed that differential mortality between life stages can stabilize or destabilize the evolutionary equilibrium where the population experiences disruptive selection, we next investigate the cause of this phenomenon. We do so by examining the effects of differential mortality between life stages on the ecological dynamics. This analysis reveals that the population reaches a larger size when mortality is biased towards juveniles than towards adults, regardless of the feeding niche trait (figure 3A). This is because differential mortality between life stages alters the population composition. When mortality is more biased towards juveniles, the fraction of adults, which are the only individuals that effectively contribute to reproduction, increases. As a result, the population reproduction rate rises (figure 3B). With a larger population, the food resources are depleted to a lower level (figure 3C) and competition thus intensifies. Strong competition induces frequency-dependent selection, which is necessary for evolution to drive a population towards a local minimum of the fitness landscape^8^. On the contrary, weak intraspecific competition, that occurs when mortality is biased towards adults, results in frequency-independent selection, precluding the possibility of diversification. This effect is robust over a broad range of maximum stage-specific mortality (figure S1).

### Juvenile-biased mortality reduces the constrains for diversification

For diversification to occur, productivity must be high. Indeed, an analysis of the fitness gradient experienced by a population colonizing an environment with two different food resources indicates that low productivity prevents diversification (figure 4A, 4B). There exists therefore a threshold minimum of productivity that enables diversification, or in other words, a minimum level of productivity enabling the local fitness minimum (trait value 1.5) to be an attractor of the evolutionary dynamics. This threshold is strongly influenced by differential mortality between life stages. As mortality becomes more biased towards juveniles, the threshold minimum of productivity that enables diversification is shifted to a lower level of productivity (figure 4C). As a consequence, diversification can occur over a larger range of productivities, and thus is less restricted, when mortality is biased towards juveniles than when it is equal for all individuals. Juvenile-biased mortality therefore facilitates diversification. This effect is the consequence of the influence that differential mortality between life stages has on the population composition and thus in the strength of intraspecific competition. Juvenile-biased mortality by enabling strong competition relaxes the productivity requirement to induce frequency-dependent selection and hence diversification. Further analyses show that this result is robust to an uneven partitioning of productivity between the resources (figure S2).

## Discussion

Existing theory on ecological diversification has largely neglected the fact that ecological interactions do not remain constant throughout the lifetime of an organism, and that generally, small young individuals have higher mortality rates than large adults. Our results reveal that this difference in mortality between life stages is a crucial factor for ecological diversification. Indeed, we show that juvenile-biased mortality facilitates diversification, whereas adult-biased mortality constrains it. Juvenile-biased mortality is widespread in natural populations. The general rule across animal taxa is that larger organisms have higher survival than smaller conspecifics^19,20,29,30,21–28^. Our results hence suggest that these differences in survival observed in natural populations set a scenario that, in the presence of disruptive selection, facilitates competition-driven diversification. We furthermore show that this facilitating effect is the consequence of strong intraspecific competition caused by juvenile-biased mortality. Our findings show that differential mortality between life stages can modulate the strength of intraspecific competition. Although multiple effects of differential mortality between life stages on eco-evolutionary processes have been recently uncovered (e.g. ref.^14–18^), to our knowledge, its modulating effect on intraspecific competition has not been documented. Given the prominence of intraspecific competition in several ecological and evolutionary processes, this modulating effect may have important consequences beyond ecological diversification that need further research.

The model in this study is based on the assumption that individuals reproduce asexually. In sexual populations, diversification is hindered by random mating. Diversification with sexual reproduction can only occur if gene flow is reduced through the evolution of assortative mating. Although in this study, we do not investigate the evolution of assortative mating, it is very likely to evolve in our system. In fact, multiple studies have shown that if the ecological conditions for diversification are satisfied, selection favors assortative mating^11,12^, even when taking into account various developmental stages^13^.

Our findings reveal that differential mortality between life stages has a strong influence on ecological diversification. We show that juvenile-biased mortality, which is commonly observed in natural populations, enables ecological diversification over a larger range of resource productivities than if mortality is equal for all individuals. Diversification may therefore be less restricted than thought based on existing diversification models^6,7^, which neglect life stages and thus implicitly assume mortality to be equal for all individuals. Interestingly, recent work showed that juvenile-biased mortality has a stabilizing effect in complex communities, and hence is crucial for the persistence of diverse ecological communities^17^. Taken all together, these findings and the present results suggest that differential mortality between life stages plays a central role in the processes underlying the origin of new species and their persistence. If we are to understand the origin and maintenance of biodiversity, it will prove necessary to gain further insight into the ecological and evolutionary consequences of individual variability arising from developmental changes.

## Notes

### Competing Interest Statement

The authors have declared no competing interest.

