## Supporting information for "Differential stage-specific mortality as a mechanism for diversification"

This supplementary note includes:

- Supplementary text:
  1. Extended model including multiple resources, stage-specific feeding rates and metabolic maintenance
  2. Derivation of the canonical equation and curvature of the fitness landscape
- Figures S1 and S2

**SI 1. Extended model including multiple resources, stage-specific feeding rates and metabolic maintenance**

In the following, we present an extended version of the model in the main text that considers multiple resources, differences in feeding rate between life stages as well as metabolic maintenance costs. The model considers $n$ food resources with density $F_i$ ($i = 1, \ldots, n$) and $m$ emerging ecomorphs ($j = 1, \ldots, m$). We assume that there exists an optimal trait value $\theta_i$ to consume each resource. These optimal traits are ordered along a one-dimensional ecological trait space (i.e. $\theta_1 < \theta_2 < \cdots < \theta_n$) and equally distant from one another by a distance $D$.

The feeding rate of juveniles $J_j$ of the $j$th ecomorph is

$$c_{J,j}(\eta_j, \mathrm{F}) = \gamma \sum_{i=1}^{n} a_i(\eta_j) F_i$$

and of adults $A_j$ is

$$c_{A,j}(\eta_j, \mathrm{F}) = (2 - \gamma) \sum_{i=1}^{n} a_i(\eta_j) F_i.$$

Juveniles mature at a rate

$$m_{J,j}(\eta_j, \mathrm{F}) = \max\left(\varepsilon c_{J,j} - \nu, 0\right)$$

And adults reproduce at a rate

$$b_{A,j}(\eta_j, \mathrm{F}) = \max\left(\varepsilon c_{A,j} - \nu, 0\right);$$

where $\mathrm{F} = (F_1 \ldots F_i)$, and $\nu$ is metabolic maintenance cost. Analogous to the factor $\phi$, the factor $\gamma$, ranging between 0 and 2, determines the difference in feeding rate between the life stages. When $\gamma = 1$, juveniles and adults feed at the same rate. Conversely, when $\gamma > 1$ ($\gamma < 1$), juveniles feed at a faster (slower) rate than adults. The functions max implies that maturation and reproduction only take place when the ingested food exceeds the metabolic maintenance cost.

The resources dynamics follow:

$$\frac{dF_i}{dt} = \rho(F_{i\,max} - F_i) - \gamma \sum_{j=1}^{m} a_i(\eta_j) F_i J_j - (2 - \gamma) \sum_{j=1}^{m} a_i(\eta_j) F_i A_j, \qquad \text{(SI1.1)}$$

and the ecomorph population dynamics:

$$\frac{dJ_j}{dt} = b_{A,j}A_j - m_{J,j}\,J_j - (\phi\delta_s + \delta_b)\,J_j$$

(SI1.2)

$$\frac{dA_j}{dt} = m_{J,j}\,J_j - \big((2-\phi)\delta_s + \delta_b\big)A_j$$

(SI1.3)

When, $\gamma = 1$ and $\nu = 0$, this model extension is equivalent to the model presented in the main text.

For this extended model version, we derive the canonical equation of adaptive dynamics that describe the evolutionary dynamics:

$$\frac{d\eta_j}{dt} = \frac{T_{f,j}\big(\eta_j\big)N_j^*\big(\eta_j\big)\mu}{T_{s,j}\big(\eta_j\big)\,\sigma^2}\cdot\frac{1}{T_{f,j}}\cdot\frac{\partial\left(\log\left(R_0\big(\eta'_j,\eta\big)\right)\right)}{\partial\eta'_j}\Bigg|_{\eta'_j=\eta_j} = \frac{N_j^*\big(\eta_j\big)\mu}{T_{s,j}\big(\eta_j\big)\,\sigma^2}\cdot\frac{1}{R_0\big(\eta'_j,\eta\big)}\cdot\frac{\partial\,R_0\big(\eta'_j,\eta\big)}{\partial\eta'_j}\Bigg|_{\eta'_j=\eta_j}.$$

(SI1.4)

where $T_{s,j}$, replaced by the notation $T_j$ for simplicity, is the sum of the expected lifespan of an individual as a juvenile and as an adult at ecological equilibrium:

$$T_j\big(\eta_j\big) = \frac{1-\frac{m_{J,j}}{m_{J,j}+\phi\delta_s+\delta_b}}{\phi\delta_s+\delta_b} + \frac{\frac{m_{J,j}}{m_{J,j}+\phi\delta_s+\delta_b}}{(2-\phi)\delta_s+\delta_b} = m_{J,j}+\phi\delta_s+\delta_b+\frac{m_{J,j}}{\big(\delta_s(2-\phi)+\delta_b\big)\big(m_{J,j}+\phi\delta_s+\delta_b\big)}.$$

In the following section, we analytically derive an expression for $\frac{\partial R_0\left(\eta'_j,\eta\right)}{\partial\eta'_j}$.

The lifetime reproductive output of an individual, $R_0$, equals the product of the expected number of offspring produced over the lifetime of an adult and the probability of surviving until adulthood. The average lifetime of an adult is $1/[\delta_s(2-\phi)+\delta_b]$, and hence its expected offspring equals $b_{A,j}/[\delta_s(2-\phi)+\delta_b]$, where $b_{A,j} = \varepsilon\sum_{i=1}^{n} a_i\big(\eta_j\big)F_i$ . The probability of surviving until adulthood equals the number of individuals that mature, $m_{J,j}\,J_j$, divided by the number of newborns, $b_{A,j}A_j$. In the ecological equilibrium, the ratio $J_j/A_j$ can be derived from eq. 1, which is equivalent to:

$$0 = b_{A,j}A_j - m_{J,j}\,J_j - (\phi\delta_s + \delta_b)\,J_j$$

$$\frac{J_j}{A_j} = \frac{b_{A,j}}{m_{J,j}+\phi\delta_s+\delta_b}.$$

Therefore,

$$R_0 = \frac{b_{A,j}}{\delta_s(2-\phi)+\delta_b} \cdot \frac{m_{J,j}\, b_{A,j}}{b_{A,j}\left(m_{J,j}+\phi\delta_s+\delta_b\right)} = \frac{b_{A,j}\, m_{J,j}}{(\delta_s(2-\phi)+\delta_b)\left(m_{J,j}+\phi\delta_s+\delta_b\right)}.$$

(SI1.5)

The lifetime reproductive output of a mutant with trait $\eta'_j$ in the environment of the resident ecomophs with trait values $\eta$ is thus given by:

$$R_0\left(\eta'_j,\eta\right) = \frac{b_{A,j}\left(\eta'_j,\mathrm{F}(\eta)\right) \cdot m_{J,j}\left(\eta'_j,\mathrm{F}(\eta)\right)}{(\delta_s(2-\phi)+\delta_b)\left(m_{J,j}\left(\eta'_j,\mathrm{F}(\eta)\right)+\phi\delta_s+\delta_b\right)}.$$

(SI1.6)

Defining

$$\lambda(\eta'_j,\eta) = \sum_{i=1}^{n} a_i\left(\eta'_j\right)F_i\,(\eta)$$

$$C_m = \varepsilon\gamma,\ C_b = \varepsilon(2-\gamma)$$

$$C_1 = \phi\delta_s+\delta_b,\ C_2 = \delta_s(2-\phi)+\delta_b,$$

eq. SI1.6 can be rewritten as follows

$$R_0\left(\lambda(\eta'_j,\eta)\right) = \frac{\left(\,C_b\lambda\left(\eta'_j,\eta\right)-\nu\,\right)\left(\,C_m\lambda\left(\eta'_j,\eta\right)-\nu\,\right)}{C_2\left(C_m\lambda\left(\eta'_j,\eta\right)+C_1\right)}.$$

$\frac{\partial R_0\left(\eta'_j,\eta\right)}{\partial \eta'_j}$ can be reformulated as:

$$\frac{\partial R_0\left(\eta'_j,\eta\right)}{\partial \eta'_j} = \frac{\partial R_0(\lambda)}{\partial \lambda} \cdot \frac{\partial \lambda\left(\eta'_j,\eta\right)}{\partial \eta'_j},$$

(SI1.7)

where

$$\frac{\partial R_0(\lambda)}{\partial \lambda} = \frac{C_b {C_m}^2\lambda^2 + 2\,C_b\,C_m\,C_1\lambda - \,C_1\nu(\,C_m+\,C_b) - C_m\nu^2}{C_2(C_m\lambda+C_1)^2}$$

and

$$\frac{\partial\lambda(\eta'_j,\eta)}{\partial\eta'_j} = \sum_{i=1}^{n} \frac{\alpha\,(\theta_i - \eta'_j)}{\tau^2} \exp\left[\frac{-(\theta_i - \eta'_j)^2}{2\tau^2}\right] F_i(\eta).$$

**SI 2: Derivation of the canonical equation and curvature of the fitness landscape**

Following Durinx et al. 2008[1], the canonical equation of adaptive dynamics, that determines the rate of change of the trait, is given by:

$$\frac{d\eta_j}{dt} = \frac{T_{f,j}(\eta_j)\, N_j^*(\eta_j)\mu}{T_{s,j}(\eta_j)\ \sigma^2} \cdot \left.\frac{\partial s(\eta_j', \eta)}{\partial \eta_j'}\right|_{\eta_j' = \eta_j}, \tag{SI2.1}$$

where $T_{f,j}(\eta_j)$ is the average age at giving birth of an individual of the $j$th ecomorph, $T_{s,j}(\eta_j)$ is its expected life span, $N_j^*(\eta_j)$ is its density at ecological equilibrium, $\mu$ is the mutation rate per birth event, $\sigma^2$ is the variance of the trait offspring distribution, and $\left.\frac{\partial s(\eta_j', \eta)}{\partial \eta_j'}\right|_{\eta_j' = \eta_j}$ is the selection gradient. $s(\eta_j', \eta)$ is the long-term population growth of a mutant type $\eta_j'$ in an environment that is dominated and hence determined by a resident (potentially polymorphic) population with trait values $\eta$, and it is related to the $R_0(\eta_j', \eta)$ following:

$$s(\eta_j', \eta) = \frac{\log\left(R_0(\eta_j', \eta)\right)}{T_{f,j}(\eta_j)}.$$

Therefore, the rate of change of the feeding niche trait can be rewritten as follows:

$$\frac{d\eta_j}{dt} = \frac{T_{f,j}(\eta_j) N_j^*(\eta_j)\mu}{T_{s,j}(\eta_j)\ \sigma^2} \cdot \frac{1}{T_{f,j}} \cdot \left.\frac{\partial\left(\log\left(R_0(\eta_j', \eta)\right)\right)}{\partial \eta_j'}\right|_{\eta_j' = \eta_j} = \frac{N_j^*(\eta_j)\mu}{T_{s,j}(\eta_j)\ \sigma^2} \cdot \frac{1}{R_0(\eta_j', \eta)} \cdot \left.\frac{\partial\, R_0(\eta_j', \eta)}{\partial \eta_j'}\right|_{\eta_j' = \eta_j}. \tag{SI2.2}$$

In the main text $T_{s,j}$ was replaced by the notation $T_j$ for simplicity, and it is sum of the expected lifespan of an individual as a juvenile and as an adult at ecological equilibrium:

$$T_j(\eta_j) = \frac{1 - \frac{m_{J,j}}{m_{J,j} + \phi\delta_s + \delta_b}}{\phi\delta_s + \delta_b} + \frac{\frac{m_{J,j}}{m_{J,j} + \phi\delta_s + \delta_b}}{(2-\phi)\delta_s + \delta_b} = m_{J,j} + \phi\delta_s + \delta_b + \frac{m_{J,j}}{(\delta_s(2-\phi) + \delta_b)(m_{J,j} + \phi\delta_s + \delta_b)},$$

where $m_{J,j} = \varepsilon \sum_{i=1}^{n} a_i(\eta_j) F_i$.

In the following section, we analytically derive an expression for $\frac{\partial R_0(\eta_j', \eta)}{\partial \eta_j'}$.

The lifetime reproductive output of an individual, $R_0$, equals the product of the expected number of offspring produced over the lifetime of an adult and the probability of surviving until adulthood. The average lifetime of an adult is $1/[\delta_s(2-\phi)+\delta_b]$, and hence its expected offspring equals $b_{A,j}/[\delta_s(2-\phi)+\delta_b]$, where $b_{A,j} = \varepsilon\sum_{i=1}^{n} a_i(\eta_j)F_i$ . The probability of surviving until adulthood equals the number of individuals that mature, $m_{J,j}\, J_j$, divided by the number of newborns, $b_{A,j}A_j$. In the ecological equilibrium, the ratio $J_j/A_j$ can be derived from eq. 1, which is equivalent to:

$$0 = b_{A,j}A_j - m_{J,j}\, J_j - (\phi\delta_s + \delta_b)\, J_j$$

$$\frac{J_j}{A_j} = \frac{b_{A,j}}{m_{J,j} + \phi\delta_s + \delta_b}.$$

Therefore,

$$R_0 = \frac{b_{A,j}}{\delta_s(2-\phi)+\delta_b}\cdot\frac{m_{J,j}\, b_{A,j}}{b_{A,j}\left(m_{J,j}+\phi\delta_s+\delta_b\right)} = \frac{b_{A,j}\, m_{J,j}}{(\delta_s(2-\phi)+\delta_b)\left(m_{J,j}+\phi\delta_s+\delta_b\right)}.$$

(SI2.3)

The lifetime reproductive output of a mutant with trait $\eta'_j$ in the environment of the resident ecomophs with trait values $\eta$ is thus given by:

$$R_0\left(\eta'_j,\eta\right) = \frac{b_{A,j}\left(\eta'_j, \mathrm{F}(\eta)\right)\cdot m_{J,j}\left(\eta'_j, \mathrm{F}(\eta)\right)}{(\delta_s(2-\phi)+\delta_b)\left(m_{J,j}\left(\eta'_j, \mathrm{F}(\eta)\right)+\phi\delta_s+\delta_b\right)}.$$

(SI2.4)

Defining

$$\lambda(\eta'_j,\eta) = \sum_{i=1}^{n} a_i\left(\eta'_j\right)F_i\,(\eta)$$

$$C_1 = \phi\delta_s + \delta_b,\ C_2 = \delta_s(2-\phi)+\delta_b,$$

eq. SI2.4 can be rewritten as follows

$$R_0\left(\lambda(\eta'_j,\eta)\right) = \frac{\left(\varepsilon\lambda\left(\eta'_j,\eta\right)\right)^2}{C_2\left(C_m\lambda\left(\eta'_j,\eta\right)+C_1\right)}.$$

$\frac{\partial R_0\left(\eta'_j,\eta\right)}{\partial\eta'_j}$ can be reformulated as:

$$\frac{\partial R_0(\eta'_j,\eta)}{\partial \eta'_j} = \frac{\partial R_0(\lambda)}{\partial \lambda} \cdot \frac{\partial \lambda(\eta'_j,\eta)}{\partial \eta'_j},$$

(SI2.5)

where

$$\frac{\partial R_0(\lambda)}{\partial \lambda} = \frac{\varepsilon^3\lambda^2 + 2\varepsilon^2 C_1\lambda}{C_2(\varepsilon\lambda + C_1)^2}$$

and

$$\frac{\partial \lambda(\eta'_j,\eta)}{\partial \eta'_j} = \sum_{i=1}^{n} \frac{\alpha\,(\theta_i - \eta'_j)}{\tau^2} \exp\left[\frac{-(\theta_i - \eta'_j)^2}{2\tau^2}\right] F_i(\eta).$$

Directional selection halts where the selection gradient vanishes, in other words when eq SI2.5 equals 0. At this point, selection can be stabilizing if this trait value corresponds to a fitness maximum, or disruptive if it corresponds to a fitness minimum. The curvature of the fitness function therefore determines whether evolution halts or an ecomorph splits into two different ecomorphs. Hence, the second derivative of the fitness function with respect to $\eta'_j$ evaluated at the ecological equilibrium when ecomorphs have traits $\eta$ allows us to determine whether a diversification event occurs.

$$\frac{\partial^2 R_0(\eta'_j,\eta)}{\partial {\eta'_j}^2} = \frac{\partial^2 R_0(\lambda)}{\partial \lambda^2} \cdot \left(\frac{\partial \lambda(\eta'_j,\eta)}{\partial \eta'_j}\right)^2 + \frac{\partial R_0(\lambda)}{\partial \lambda} \cdot \frac{\partial^2 \lambda(\eta'_j,\eta)}{{\eta'_j}^2},$$

(SI2.6)

where the two unknown terms are:

$$\frac{\partial^2 R_0(\lambda)}{\partial \lambda^2} = \frac{2\varepsilon^2 {C_1}^2}{C_2(\varepsilon\lambda + C_1)^3}$$

$$\frac{\partial^2 \lambda(\eta'_j,\eta)}{{\eta'_j}^2} = \sum_{i=1}^{n} \frac{\alpha\,\left((\theta_i - \eta'_j)^2 - \tau^2\right)}{\tau^4} \exp\left[\frac{-(\theta_i - \eta'_j)^2}{2\tau^2}\right] F_i(\eta).$$

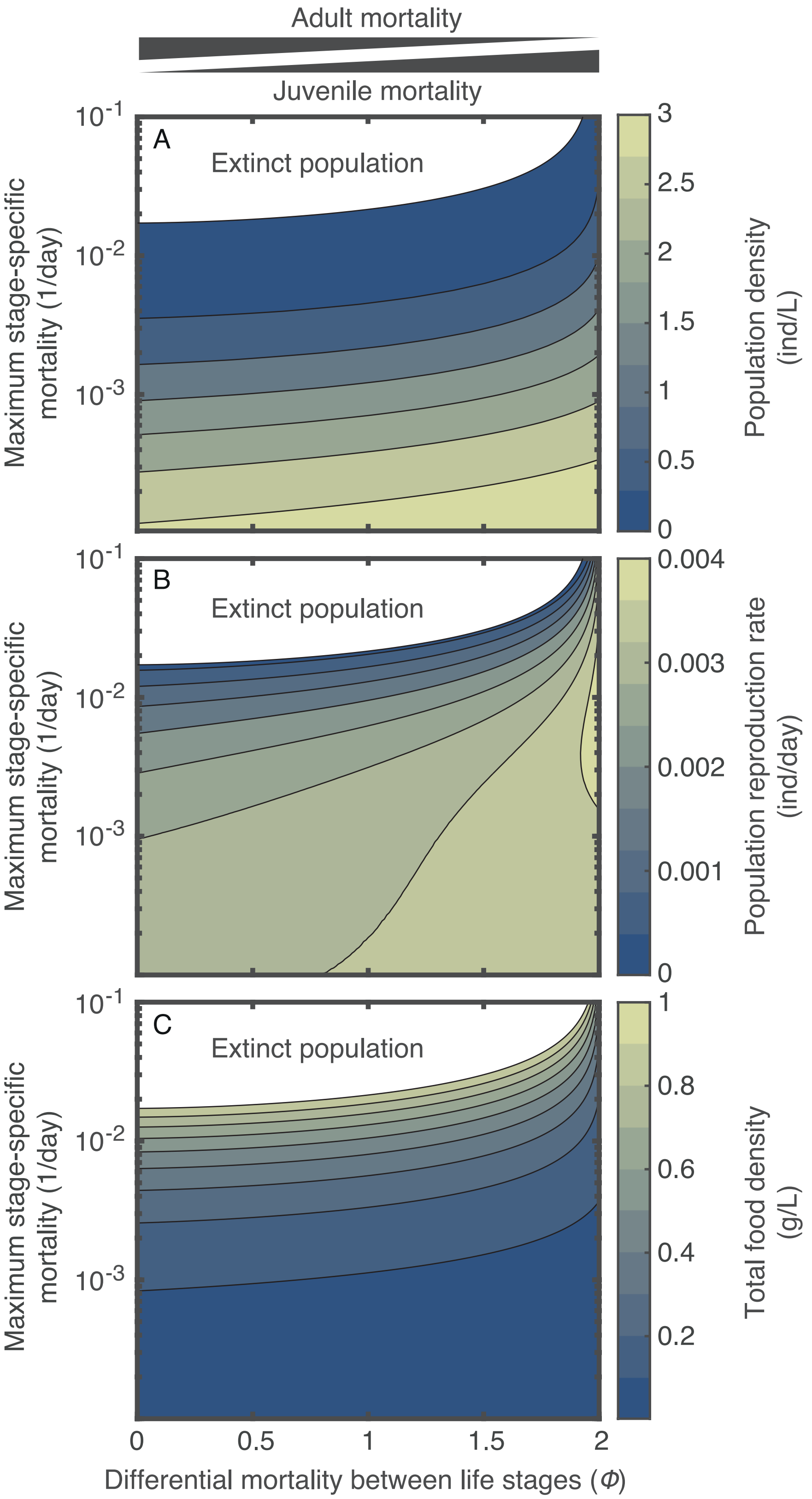


Figure S1. A) Population density, B) population reproduction rate and C) total food density in the ecological equilibrium as a function of the differential mortality between life stages and the maximum stage-specific mortality. The population density is the sum of the juvenile and adult density. The total food density is the sum of the densities of the two food resources. As in figure 3, no evolutionary dynamics are considered. The feeding niche trait is fixed and equal to 1.4. The population is extinct in the white region. For every maximum stage-specific mortality, the population density and population reproduction rate increase with increasing differential mortalities between life stages, whereas the total food density decreases. Parameter values as in table 1.

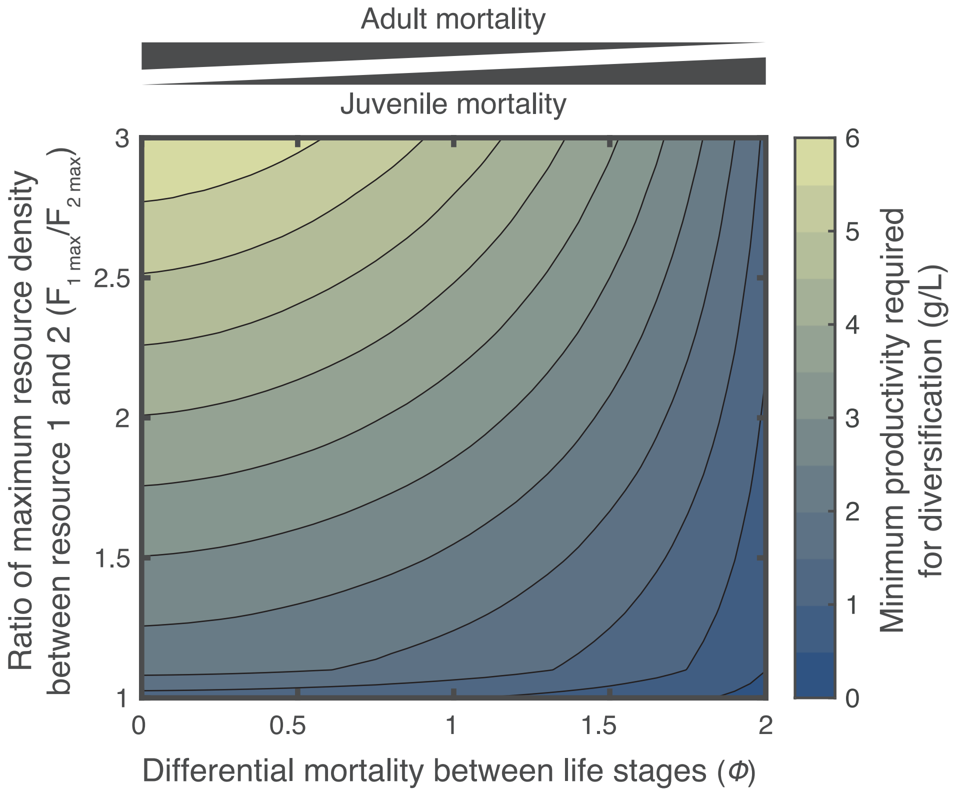


Figure S2. Minimum productivity required for diversification as a function of differential mortality between life stages (horizontal axis) and the ratio between the maximum resource density of the resources (vertical axis). For every ratio of maximum resource density between the resources, the minimum productivity required for diversification decreases with increasing bias of mortality towards the juvenile stage. The range of the ratio is varied between 1 and 3; when the ratio is 1, the maximum resource densities are equal (as in figure 4), when the ratio is larger than 3.2, the population evolves towards the vecinity of the optimum to feed on resource 1 and selection is balancing. The values of the ratio between 1 and 3 therefore cover most of the range over which disruptive selection can occur. Parameter values as in table 1.
